# Preparation of Organ Scaffolds by Decellularization of Pancreas and Re-functionalization

**DOI:** 10.1101/513465

**Authors:** K Uday Chandrika, Rekha Tripathi, T Avinash Raj, N. Sairam, Vasundhara Kamineni Parliker, VB Swami, Nandini Rangaraj, J Mahesh Kumar, Shashi Singh

## Abstract

Extracellular matrix of each tissue is unique in composition, architecture and finer details that support the very identity of the organ by regulating the status/character of the cells within it. Tissue engineering centers around creating a niche similar to the natural one, with a purpose of developing an organ/oid. In this study, whole organ decellularization of pancreas was attempted followed by reseeding it with adult mesenchymal stem cells. Decellularization completely removes cells leaving behind extracellular matrix rich scaffold. After reseeding, mesenchymal stem cells differentiate into pancreas specific cells. Upon transplantation of recellularized pancreas in streptozotocin induced diabetic mice, this organ was capable of restoring its histomorphology and normal functioning. Restoration of endocrine islets, the exocrine acinar region, and vascular network was seen in transplanted pancreas. The entire process of functional recovery took about 20 days when the mice demonstrated glucoregulation, though none achieved gluconormalization. Transplanted mice upon feeding show insulin and c-peptide in circulation. This process demonstrates that natural scaffolds of soft organs can be refunctionalized using recipients cells to counter immune problems arising due to organ transplantation.

## 1. INTRODUCTION

Transplantation is the only hope for scores of patients with irreversible or end organ failure. Organ supply remains the first hurdle, followed by the immune-complications and rejection failures. The reason behind the hype of tissue engineering has been its potential to manage the debilitating diseases involving organ damage/failures. The usual approach of tissue engineering has been to create a construct that would mimic an organ functionally to attenuate and ultimately overcome the dysfunctionality.

There are innumerous approaches with variety of materials (synthetic, natural, combinations) and manifold techniques to create scaffolds that can be seeded with cells to create a biomimetic tissue constructs (Karp et al., 2007, Langer and Tirrell, 2004). Most of these have shown partial success. Attempts have been made to derive scaffolds from the whole organs/tissues by decellularization (Scarrit et al., 2015, Spector, 2016) and such attempts for hard tissue like trachea, or bladder etc. have been successful in clinical trials (Baiguera and Macchiarini, 2010, Hinderer and Schenke-Layland, 2013, Macchiarini et al., 2008). Decellularization is a process by which organs are denuded of cells, leaving behind the entire blueprint in form of the architectural scaffold and cues for cell preservation and homeostasis. With the final goal to form a remodeled functional organ, the extracellular matrix obtained with close to ‘intact’ details and composition is seeded with cells. Successful recellularization of organ scaffolds has been reported in liver (Baptista et al., 2011,); lungs (Peterson et al., 2010, Song et al., 2011); nerves (Crapo et al., 2012); tendons (Martinello et al., 2013); bladder (Loai et al., 2010), pancreas (Goh et al., 2013; Peloso et al., 2016), heart and valves (Ott et al., 2008; Zhou et al., 2010); kidney (Song et al., 2013, Schmitt et al. 2017).

Tissue decellularization has been attempted by flushing the organ through vascular channels with mild detergents, enzymatic solutions, chemicals or mechanical, or a combination of few of these (Crapo et al., 2011, Peloso et al., 2015). A fine balance is sought between preserving the tissue architecture with essential structural proteins, growth stimulating signals/factors and complete removal of cellular, DNA or any immunogenic content. Various reagents like SDS, Triton X, per-acetic acid, EDTA and trypsin have been reported in various studies for different organ decellularization (Fu et al., 2014).

Diabetes remains one of the most inadequately managed disease; with the population ending up with morbidity and mortality due to hyperglycemia related organ damage. Long term benefits are sought with whole pancreas transplantations or islets but as usual poor availability and use of immune suppressants are some of the limitations (Friedman and Friedman, 2002, Qi, 2014). Islets are encased in semisynthetic, semipermeable membrane to avoid immune surveillance of host but cell survival and functionality still remains an issue (Qi, 2014). Addition of ECM components affects cell survival and insulin release and also forms the background for using the decellularized ECM for refunctionalization (De Carlo et al., 2010, Lucas Clerc et al., 1993, Mirmalek-Sani et al., 2013, Salvay et al., 2008).

Managing a disease with stem cells has been priority in regenerative medicine. For therapeutics, stem cells from embryonic sources or iPS (induced pluripotent stem cells) are debatable as these cells are highly tumorogenic (Herberts et al., 2011). Mesenchymal stem cells available from a variety of sources; are one class of cells that have been on the radar for their potential clinical applications. These can be obtained in a large number without much manipulation and have been used in a few clinical applications (Ikebe and Suzuki, 2014, Squillaro et al., 2015, Thomsen et al., 2014). Apart from the immune advantage these cells ‘may’ offer actual regenerative advantage (Squillaro et al., 2015, Thejaswi et al., 2015). One of the problems with these cells is their diminishing proliferating potential in culture conditions due to ageing. Once injected at the site or systemically, actual implantation at the site has been quite low. Owing to these, long term benefits of these procedures have been questionable. In this study, we have explored the possibility of use of MSCs from murine or human sources to repopulate the scaffold obtained by decellularization and functionalize the organ.

Use of MSC was also necessitated as organ like pancreas is developmentally made up with distinctive combination of cell lineages derived from two distinct endodermal buds (Jennings et al., 2015). The exocrine portion is made up of acinar cells that secrete digestive juices into a ductal system draining into intestines. Multiple cell types comprise the islets that form the endocrine system of pancreas. Progenitors or differentiated cells are one option that is quite popular for reseeding (Scarritt et al., 2015) but one needs multiple cell types to be introduced. Repopulating with naive cells made more sense than using single or many differentiated lineages. Second this system offers us the advantage of studying development and differentiation of these cells. Stem cells in an inductive environment like ECM would adopt the appropriate pathway of differentiation.

In this paper, we present our study dealing with decellularization to create an acellular scaffold, reseeding with mesenchymal stem cells for recellularization of pancreas. Implantation of this construct restores the glucoregulation in a streptozotocin induced diabetic mice.

## 2. Materials and Methods

### 2.1 Mouse pancreas harvest and perfusion decellularization

Balb/c mice about 6-8 weeks of age were used for the experiment. The animals were maintained at standard environmental conditions (temperature 22–25° C, humidity 40–70 % with 12:12 dark/light photoperiod) approved by the Committee for the Purpose of Control and Supervision of Experiments on Animals (CPCSEA) whereas all the experimental protocols were approved by Institutional Animal Ethics Committee (IAEC), CCMB.

Balb/c mice, between the ages of 6–8 weeks were euthanized by cervical dislocation. Sterile conditions were maintained during the whole procedure. The pancreas was carefully detached free from all adjacent structures including the stomach, intestine, spleen while retaining the pancreatic and spleenic vessels intact. Insulin needle of 8mm size 31G was inserted into the hepatic portal vein to enable perfusion of the organ.

The isolated pancreas with the cannulated needle was connected to a perfusion system to allow retrograde perfusion at 5ml/min. PBS was perfused in through the hepatic portal vein and throughout the vasculature of the pancreas and exited through the spleenic vein. After complete removal of blood, non-ionic detergent, 1% Triton X-100, and 0.1% Ammonia (Sigma Aldrich) in deionized water (Mirmalek-Sani et al. 2013) was used for perfusion to detach cells and cell debris from the pancreas. After the tissues became translucent (roughly around 12h), subsequent steps of perfusion were using DNase 2000U/ml in (PBS) (~4h) perfusion followed by a final washing step of with Pencillin/Streptomycin (100U/ml) in PBS. The pancreas was perfused with PBS for additional 48 hours to clear remaining cellular debris. The entire operation was carried out in a sterile chamber. The same protocol of decellularization was used for rat and rabbit pancreas.

In case of human fetal organs, the study was approved by ethical committees of the hospital and CCMB. Informed consent was obtained from the patients undergoing medical termination of Pregnancy. A total of four human fetal pancreases were used for the study. Pancreas was harvested after bringing the abortus under sterile conditions and cannulated from the hepatic portal vein. The protocol for decellularization was followed in the same manner as listed for mouse pancreas.

### 2.2 Cell culture

The human placenta derived MSC (hPL-MSC) and mouse Adipose MSC (mAD-MSC) were maintained in IMDM (Iscove’s Modified Dulbecco’s medium; Gibco USA) containing 15 % FBS with 100U/ml Pencillin-Streptomycin (Life Technologies) at 37°C and in a 95% air/5% CO2 atmosphere. MSC between passages 3–6 were used for recellularizing the tissue scaffold.

### 2.3 Recellularization

Confluent cultures of MSC were trypsinized and 6-10 × 10^5^ cells were collected in 3mL of DMEM medium. The cell suspension was introduced into the decellularized pancreas by means of perfusion via the hepatic portal vein in 3 infusion steps. Each time 1mL cell suspension containing 2 × 10^5^ cells was perfused for over 20 min. The next infusion of cells was passed after 2 hrs. Throughout the process, pancreata was continuously perfused with CMRL 1640 (GIBCO) culture medium for 5 days. Cannula to the portal vein was kept inserted to allow perfusion feeding of media continuously.

Rat, rabbit and human fetal pancreas were seeded using the same protocol using about 12-15 × 10^5^ cells in each case.

### 2.4 Induction of Diabetes in mice

Balb-C mice about 6-8 weeks of age were used for the experiment. The animals were injected with a single dose of 160mg/Kg bodyweight streptozotocin (Graham et al 2011). Glucose levels were checked by tail vein puncture with a glucometer till the animal showed higher glucose levels confirming its diabetic status; which was attained by day 5 usually. The animals were maintained at standard environmental conditions (temperature 22–25° C, humidity 40–70 % with 12:12 dark/light photoperiod).

### 2.5 *In vivo* implantation of decellularized pancreas

The diabetic mouse was anaesthetized by intraperitoneal administration of ketamine (50mg/Kg body weight) and xylazine (5 mg/Kg body weight). The surgical site was prepared in sterile fashion using 70% isopropyl alcohol followed by the placement of sterile drapes. Using a scalpel a skin incision was made near right abdominal region, the two layer of abdominal muscle were cut open to expose the pancreas. The pancreas was rinsed with normal saline and the recellularized scaffold was placed above the pancreas. Single simple interrupted suture was made at anterior end of scaffold with anterior end of pancreas and similarly at the posterior end of pancreas by using absorbable suture (3-0 prolene) material. Then the muscle was opposed with simple interrupted suture using absorbable suture material. The skin was opposed with simple interrupted suture using silk thread [6 −0; non absorbable]. Mice was administered antibiotic (enterocin 100mg/ lit of water) and analgesic (meloxicam −1mg/ kg body weight) for the period of 5 days post surgery.

The animals that survived the surgical procedure without complications were monitored for fasting blood glucose level every alternate day by bleeding through tail vein.

### 2.6 Histopathology and Immunohistochemistry (IHC)

Native, decellularized and recellularized pancreata were processed for histochemical analysis. Routine haematoxylin-eosin staining was done and sections were examined in Axioimager-2 (Zeiss, Germany). Immunofluorescence staining was performed using primary antibodies as listed in Supplementary Table 1 and counter stained with appropriate fluorescent tagged second antibody. The slides were examined in LEICA TCS SP5 with appropriate filters after counter staining with DAPI for nuclear staining. Analysis was performed using LEICA Application suite X version 2.0. 2.15022

For qualitative determination of presence of glycosaminoglycans (GAG), staining was carried out on decellularized pancreas sections. Briefly, after de-parafinization and dehydration, sections were directly stained for 5 min with Safranine O (SRL) stain at room temperature. The slides were washed in running tap water for 5 min followed by a distilled water rinse and mounted with glycerol. Slides were examined in Axioimager-2 microscope.

### 2.7 Proteomics analysis

The decellularized Balb/c mouse pancreas scaffolds were washed in PBS and homogenized-lysed for 1 hr in Tissue homozilyser II (Qiagen). The ECM proteins were dissolved in lysis buffer containing Tris HCl 0.06M, SDS 2.5% and β- mercaptoethanol 1.25%. The lysate was centrifuged and supernatant was mixed with 2x reducing SDS sample buffer and analyzed on SDS-PAGE. Each sample was reduced with 2.5mM Tris (2-carboxyethyl) phosphine and alkylated with 50mM iodoacetamide, followed by in-gel digestion with 20ng/μl of trypsin (Promega) for 16-20 h (overnight) at 37°C. Peptide digests were analyzed by nano Liquid Chromatography-tandem mass spectrometry (LC-MS/MS), on a Thermo Fisher LTQ Orbitrap Velos^TM^ connected to a Waters Acquity UPLC system (Waters Corp., Milford, MA), using a 90 minute gradient. Protein identification was performed with Proteome Discoverer 1.3 using the Sequest search engine. Database searches used the Uniprot complete mouse database (downloaded in December 2014 merged with a contaminant database from ABSciex, 50838 sequences, 24435643 residues). Settings used for protein ID mapping were for a full trypsin digest, with two missed cleavages, one static modification (cysteine carbamidomethylation), two dynamic modifications (oxidized methionines and hydroxyproline), mass tolerance of 10 ppm for precursor mass and 0.5 Da for fragment masses. Percolator; a post-processing software using a target/decoy database approach, was used to evaluate the accuracy of peptide identifications. Peptide identifications were filtered with a q-value cutoff of 0.01 (1% global False Discovery Rate, FDR). Proteins were grouped using the maximum parsimony principle and this list was imported to ProteinCenter (Thermo) and compared with the IPI mouse database for statistical analysis.

Further analysis of the protein data was carried out using online STRING Software (https://string-db.org).

### 2.8 Western blot analysis

Decellularized scaffolds were homogenized-lysed for 1 hr in Tissue homozilyser II (Qiagen). The ECM proteins were analyzed on SDS PAGE. The proteins were transferred on to a membrane (Millipore IVD HYBOND–C Extra membrane) by slot transfer under vacuum. The membrane was blocked using milk proteins and incubated overnight with primary antibodies against Collagen type I, Collagen type VI, laminin and fibronectin. After washing, the membranes were treated with HRP conjugated secondary antibodies and visualized by ECL.

### 2.9 DNA quantification

The decellularized and native pancreata (n=2) were used for total DNA isolation. After digestion with proteinase K and 0.1% SDS solution and TEN buffer at 55°C for 2 h. DNA was isolated by phenol chloroform extraction method. DNA was analyzed by agarose gel electrophoresis and the absolute amount of DNA (ng/mL) was quantified by Nanodrop spectrophotometer.

### 2.10 Real time Analysis

RNA was isolated from the grafted pancreas directly using Trizol reagent and converted to cDNA using superscript II^TM^. Glucose-6 phosphate dehydrogenase (*Gapdh*), 18S ribosomal RNA and β-2 microglobulin (B2M) served as internal controls. Real-time PCR was carried out for *PDX1*; *NGN3*, *NKX6.1* and *PPY* genes. The primers were custom synthesized by Bioserve India Ltd. and optimized using a crosswise combination matrix (Supplementary Table 2: primer sequences). A total of 25 ng cDNA was used for real-time PCR with SYBR^®^ Green as indicator using the Applied Biosystem (7900) HT fast real-time PCR system. Fold changes in gene expression were calculated by DDCT method. Statistical analysis was carried out using GraphPad prism sofware.

### 2.11 *In situ* hybridization

*In situ* hybridization was carried out on sections of implanted grafts re-cellularized with mouse adipose and human placental cells as described in Steck et al. 2010. Hybridization was carried out using digoxigenin labeled (DIG) mouse specific probes for mouse SINE/B1 and SINE/B2 repetitive elements and human specific probes for Human Specific Alu repeats. In brief, probes were prepared by using specific primers (Table 2b) using DIG labeling PCR kit (ROCHE). DIG labeled PCR products were purified using the nucleospin gel and PCR cleanup kit (Macherey ‒Nagel).

Sections were deparaffinized; rehydrated and washed three times in PBS containing 0.1% Tween at RT. After treatment with 20µg/ml proteinase K (Sigma Aldrich), sections were treated with 0.25% acetic acid containing 0.1M triethanolamine (pH 8.0) and prehybridized. Hybridization was carried out using specific probes overnight at 42°C. After washing, signals were detected using anti-DIG labeled alkaline phosphatase antibody by chromogen method using NBT/BCIP (Roche). Sections were counter stained with nuclear fast red and mounted after dehydration. Sections were examined in AxioImager II (Zeiss, Germany).

### 2.12 Estimation of Insulin and C-peptide

Blood was collected from animals used in transplantation experiments and serum was isolated. Serum was used for estimation of Insulin and c peptide using commercial kits following the procedure. Human c- peptide ELISA kit (cat # EZHCP-20K, Millipore MA), Human Insulin ELISA kit (cat # ELH- Insulin, RayBiotech GA), mouse insulin ELISA kit (cat # ELM insulin, RayBiotech GA) and rat/mouse c-peptide ELISA kit (cat # EZRMCP 2-21K, Millipore MA).

## 3. Results

### 3.1 Perfusion decellularization of mouse pancreas

Isolated mouse pancreata were cannulated and retrograde perfusion was carried out via hepatic portal vein. After flushing the organ completely of blood, the cellular contents were removed with 1% Triton X −100, a non-ionic surfactant based detergent. Triton X-100 would help in lysis of the cells resulting in removal of cell debris. The organ was later perfused with DNAse solution to rid the scaffolds of remnants of nucleic acids and finally washed free of lysate material with a thorough perfusion with buffer solution. Following perfusion, the whole pancreas turned completely translucent in about 12h (Fig. 1A-B). Histological examination by H&E staining showed complete decellularization with no traces of remnant cells. The histology showed a scaffold of fine fibrous network left behind. The scaffold sections were stained with DAPI as a quick check to look for cells and nuclear content and found to be negative. An acellular pancreas scaffold with intact gross anatomical structure of the pancreas was generated (Fig. 2 A-C). To further assess the extent of DNA removal, DNA quantification was performed using phenol-chloroform method. This method showed that DNA content decreased to about 20-50ng. The agarose gel analysis showed presence of fragmented DNA from the decellularized pancreas as opposed to high molecular weight DNA obtained from a normal un-processed pancreas (Fig. 2D). Feasibility of this perfusion decellularization technique in human-fetal pancreas, rabbit and Wistar rat were also demonstrated by following the same protocol of decellularization. At the end of the process, organ scaffold appeared translucent white in each case (Fig.1B-C). Perfusion mediated decellularization of pancreas via the vascular route efficiently removed cellular components leaving behind the relatively insoluble scaffold.

**Figure 1.**
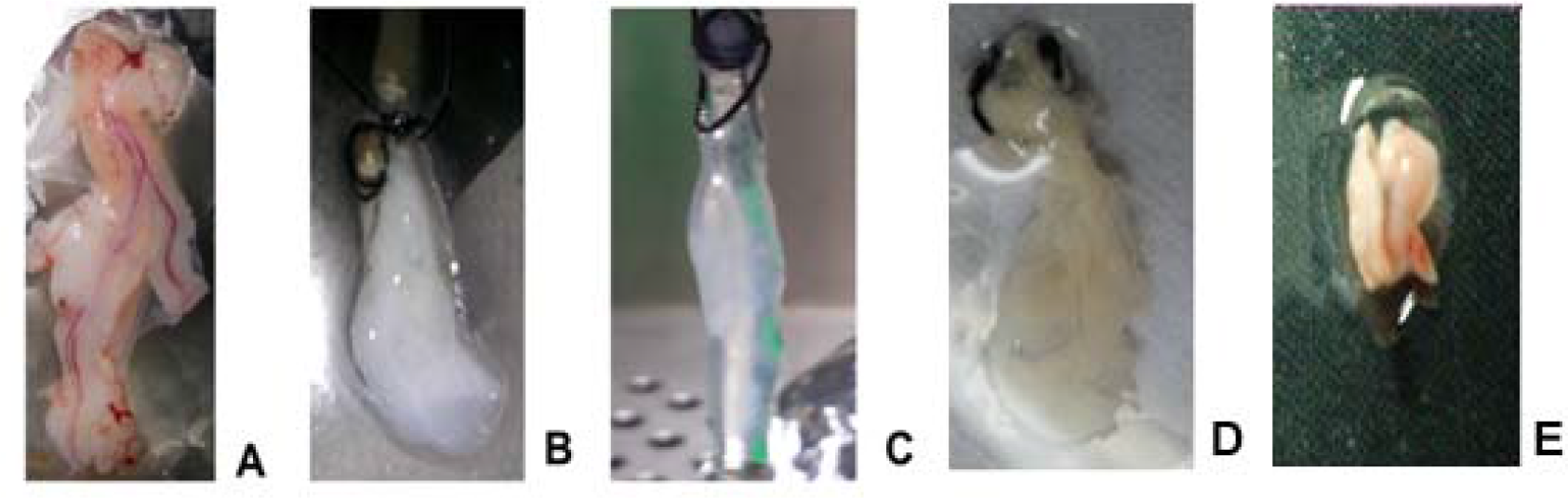
Decellularization of the pancreas. A. Whole pancreas excised from the animal. B & C. The pancreas after perfusion decellularization using detergent and enzymes. D. Histology of the decellularized pancreas showing fine fibrillar network of the extracellular matrix without cells. E. Scaffold section stained with safranin G for proteoglycans. F. Scaffold section stained with DAPI to show lack of nuclear material. F. Agarose gel picture showing degraded DNA in lane 1-4 for decellularized pancreas and Lane 5-7 Normal mouse pancreas.

**Figure 2.**
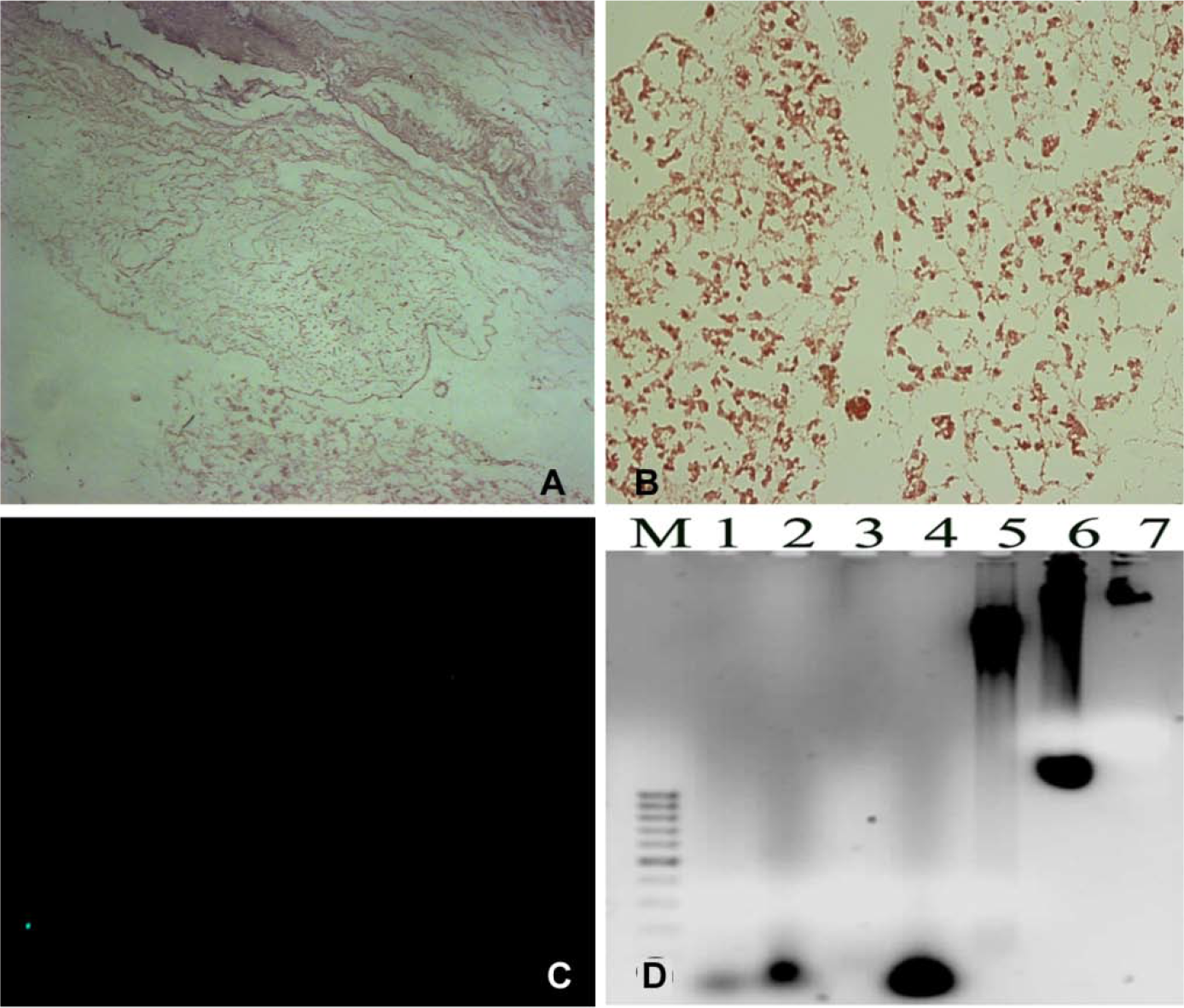
Characterization of Decellularized Scaffold. (A) Proteomic analysis of the Scaffold obtained after decellularization. Very few proteins are scored. Majority belong to the ECM (dotted set). Few belong to pancreas specific exosome proteins and few transmembrane proteins. (B) Western analysis of blotted proteins show the presence of the major ECM proteins that remain in the scaffold though there is some loss of proteins during the process of decellularization. (C) Immunochemistry of the decellularized section vs normal pancreas emphasizing the presence of ECM proteins in decellularized scaffolds.

In few trials for mouse pancreas, retrograde perfusion via pancreatic duct was also attempted for pancreas decellularization, this route did not yield a clear translucent pancreas.

### 3.2 ECM characterization: Immunohistochemistry and Mass spectrometry-based proteomics analysis

The scaffold thus obtained by decellularization was analyzed by Mass spectrometry. The histology sections were also immuno-stained for the proteins scored in proteomics. The number of proteins scored was very few (Table 1, Fig. 3A). The major proteins obtained in the scaffold were ECM proteins. Most abundant proteins scored were Collagen type I (both 1a and b) and many subtypes of collagen VI and laminin. Other ECM Proteins and glycoproteins scored were prolargin, decorin, aspirin, Dermatopontin and Desmoglein. These identities showed higher number of peptide spectral counts, accounting for their abundance.

**Table 1.**
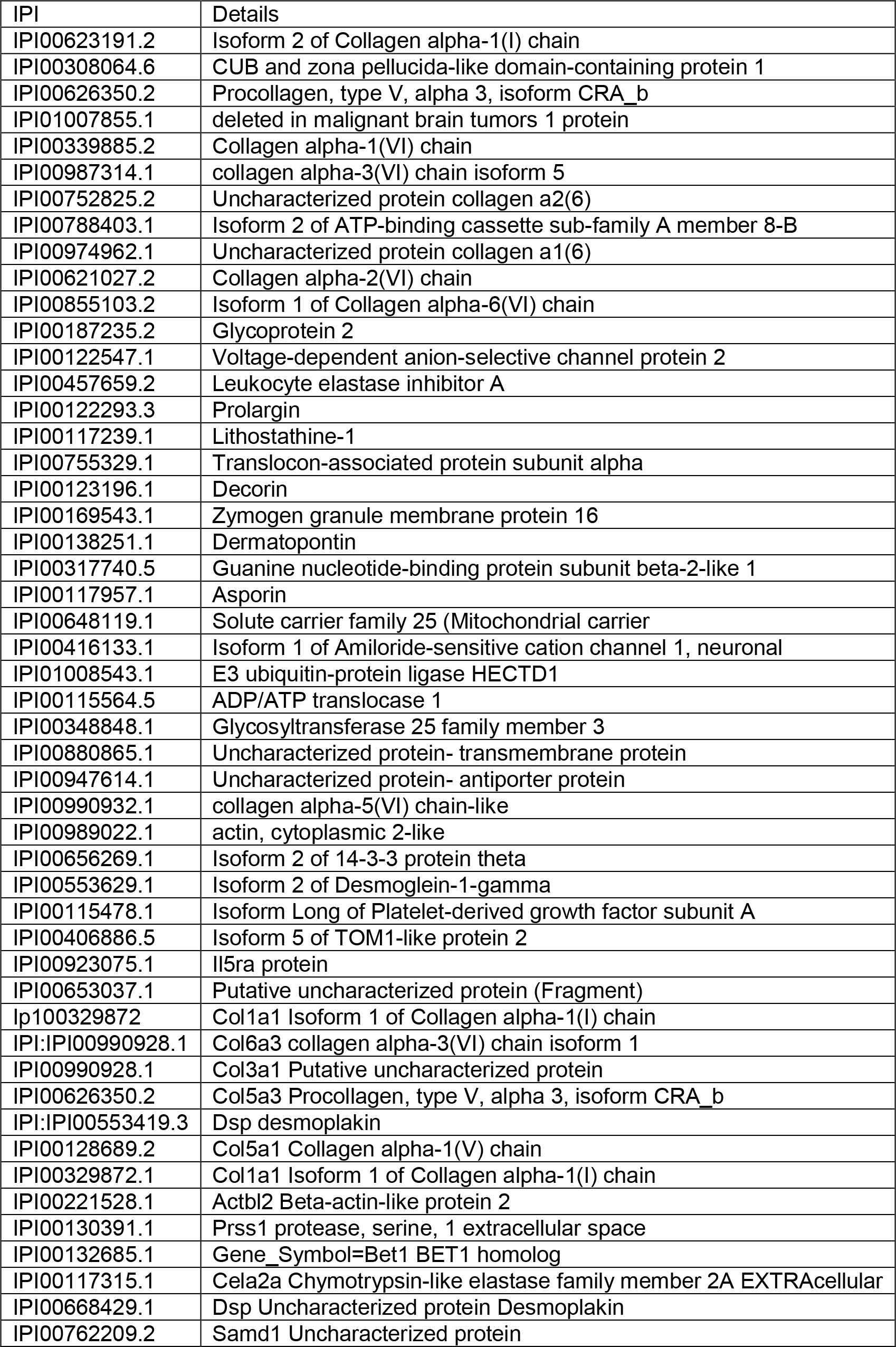
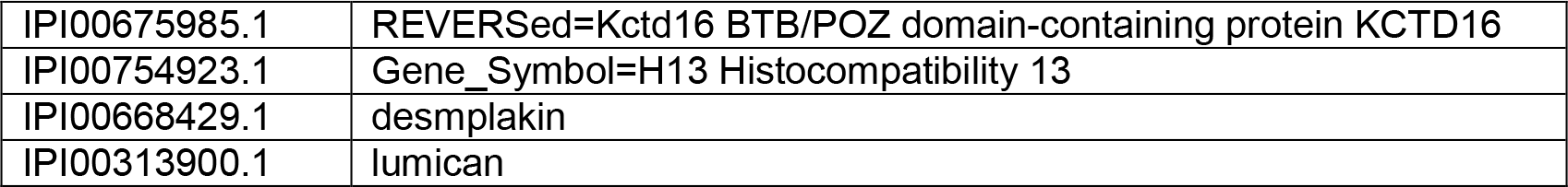
Proteomics of Decellularized scaffold of mouse Pancreas. Proteins identified in the extracellular matrix rich scaffold of pancreas obtained by Decellularization. Most of the proteins belong to ECM with few contaminants from the membranes and residues from the exocrine pancreas identifying with secretory granules.

**Table 2.** Insulin and C-peptide levels in serum of mouse implanted with recellularized pancreas. The serum collected was analyzed for human insulin and human C-peptide as well mouse insulin and mouse C peptide. – indicates values in negative or undetected.

| Model system | Human Insulin<br>U /ml | Human C<br>peptide<br>ng/ml | Mouse<br>Insulin<br>U /ml | Mouse C<br>peptide ng/ml | Blood<br>glucose<br>levels<br>mg/dl |
| --- | --- | --- | --- | --- | --- |
| Normal mice<br>n=2 | -- | 0.004+ | 29.25 +3.93 | 1.92±0.083 | 98± 1.2 |
| Diabetic mice<br>n=2 | -- | -- | 14.08 ± 1.8 | 0.031±0.01 | 341±184 |
| Mice with<br>recellularized<br>pancreas hPI-MSC n=3 | 59.64± 9.43 | 6.39 ±0.9 | -- | -- | 189 ±78 |
| Mice with<br>recellularized<br>pancreas mAD<br>n=2 | -- | 0.002+<br>0.0005 | 40.21±5.02 | 1.69 ±0.08 | 372±64 |
| Mice with<br>acellular<br>pancreas n=2 | -- | -- | 16.16 ± 3.17 | 0.22± 0.11 | 350±86 |

**Figure 3.**
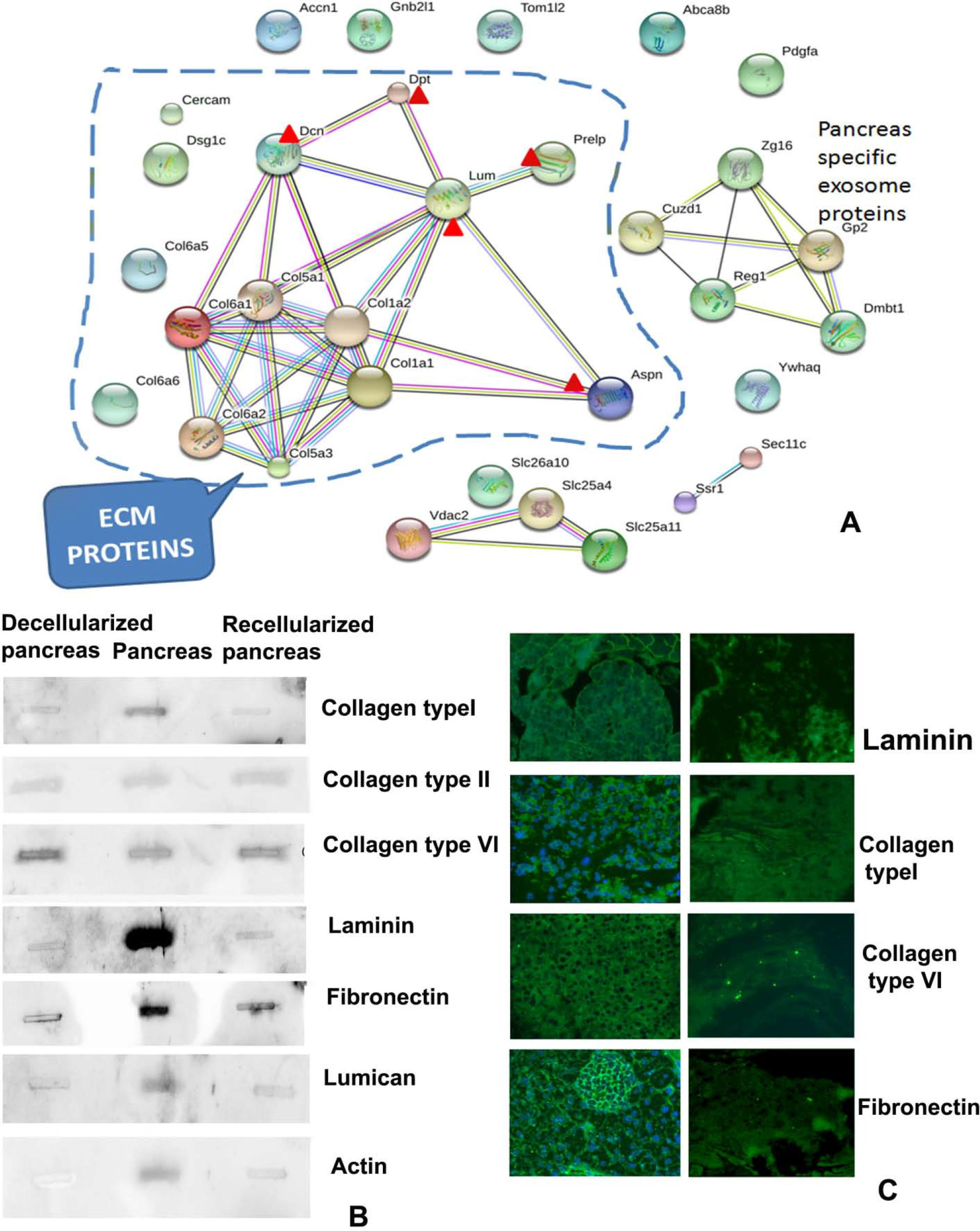
Regeneration of the pancreas. Recellularized pancreas infused with mouse adipose MSC before implantation in diabetic mouse (a). Histology of the grafted pancreas after day 15 (b) and day 40 (c) show well developed acinar cells, islets along with ducts and vasculature. (d-f) Immunohistochemistry for pancreatic markers on sections of recellularized pancreas grafted in mouse for 40 days. Insulin (Green), somatostatin (red) and glucagon (Blue) in the grafted pancreas (d). Recellularized pancreas section stained with Polypeptide y (e) and section stained for carboxypeptidase (acinar cells-red) and NGN3 (green) for beta cells (f). g -Recellularized mouse pancreas grafted in mouse excised after 40 days. h- Real time PCR analysis of beta cell markers in harvested recellularized pancreatic graft at days 7 and 30, shows enhanced expression of beta cell specific markers when compared to control i.e. MSC from placenta. j-Blood glucose profile of 6 mice grafted with recellularized pancreas show regulation of blood glucose levels. None of the mice with graft become normoglycemic but started to maintain lower glucose levels compared to diabetic mice. Red Line represents the diabetic mice with grafted recellularized pancreas and the blue lines represent the blood glucose levels of control diabetic mice.

Transporters, channel proteins, G- proteins etc. were some transmembrane proteins that were scored rarely. Similar profile was obtained for human fetal pancreas. Collagen type V and desmoplakin were also picked up in scaffold of human fetal pancreas.

Immunostaining of the decellularized pancreas stained with antibodies against some of the ECM proteins scored in proteomic analysis like Collagen type 1 and VI, fibronectin, laminin showed positive reaction (Figure 3C). All these matrix proteins showed their presence in both the normal and decellularized sections. Western analysis of the proteins obtained from the scaffolds confirmed the presence of collagen type I, VI and Laminin. Western analysis of decellularized, normal and recellularized pancreas showed there was some loss of these matrix proteins during the process of decellularization (Figure 3B). The sections were stained for Glycosaminoglycans (GAGs) using safaranin O showed a glycan rich scaffold (Figure 2 B).

### 3.3 Recellularization of scaffolds

After a thorough perfusion with PBS, extracellular matrix rich scaffold obtained was seeded with cells via the same cannula. The choice of cells was MSC from mouse adipose and human placenta that are well characterized and archived in the laboratory (Thejaswi et al. 2012). MSC were used within 3-6 passages. The organ scaffold was perfused with a total of 6-10 × 10^5^ cells in three rounds and followed with perfusion with DMEM medium for a period of 5-10 days. Histology of organs at this stage showed repopulation of the scaffold, though the histology does not appear distinctly pancreatic. Cells appear rounded and just settled (Supp fig 2a and b). Similarly with rabbit pancreas, seeding with MSC resulted in complete recovery of organ histology (Supp fig 2c).

### 3.4 Transplant experiments

The animals injected with a single dose of 160mg/Kg bodyweight streptozotocin developed high glucose levels mostly by day 4 (Figure 4i, Table 2). Streptozotocin treated diabetic mice were implanted with the recellularized pancreas intraperitoneally at a site near the host pancreas holding one end near the spleen and other near C–loop while sutured to the peritoneal muscle layer.

**Figure 4.**
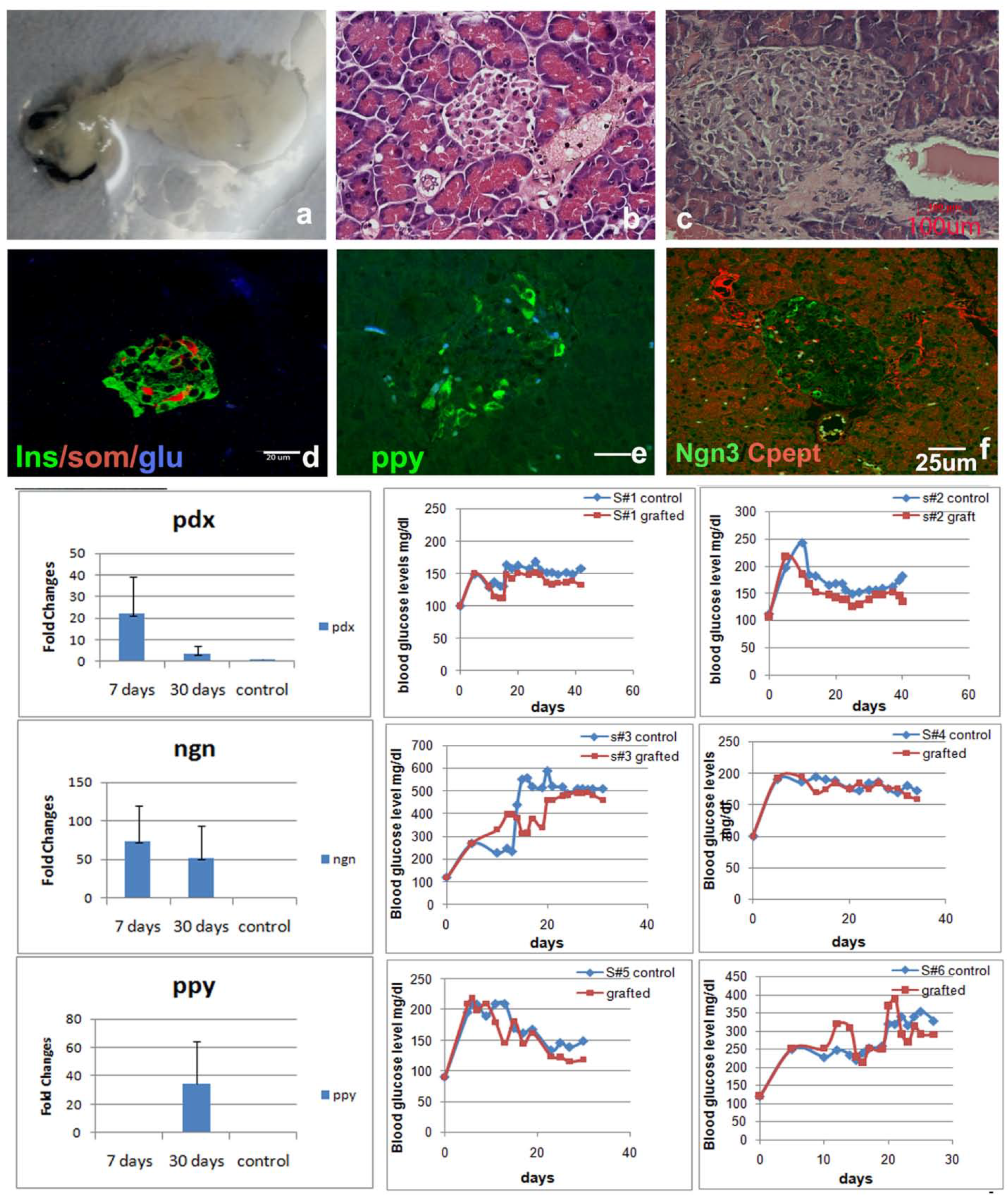
Cell tracking of human cells. Decellularized pancreas were also recellularized with human placental MSC and implanted in diabetic mice. These mice also show recovery of normal histology and tendency to achieve glucoregulation. (a) PCR products amplified using Alu primers and B1 /B2 Sine primers. Genomic DNA from mouse and human were used as template. Section of the pancreas recovered after 40 days were stained with HNA antigen and also by *in situ* hybridization using Alu (Human) and sine (Mouse) repeat sequences to show presence of human cells. b Recellularized pancreas with human cells grafted in mouse for 40 days stained for human nuclear antigen (HNA) show the presence of human cells differentiated into islets and acinar cells. The pancreas recellularized with mouse adipose MSC does not show HNA staining (c). d & e- sections of pancreas recellularized with human cells hybridized/stained with human Alu probe {inset showing nuclear staining of Alu probes} (d) and Mouse b1 and B2 sine probes (e). f & g- pancreas recellularized with mouse adipose cells stained with Human Alu probes (f) and B1& B2 mouse sine probes {inset showing nuclear staining of mouse Sine repeats probes}. Human specific Alu repeats could only be localized in human cells and there was no cross reaction.

The mice survived the surgery (12/15) and were able to recover the gluco-regulation by day 20-25 (Figure 4). The grafted pancreas developed vascular anastomoses at the graft site (S Figure 1). Pancreas of the mice recovered from these animals showed a normal histology. The islets were developed and so were the acinar cells. The vasculature was well developed and the blood cells were seen in the capillaries. Immunohistochemistry of the recellularized pancreas showed a pancreas with cells positive for endocrine markers like insulin, somatostatin, glucagon, amylin, Pancreatic poly peptide (*PPY*) in the islets and exocrine pancreas markers like carboxypeptidase in acinar cells (Figure 3a-f). qPCR analysis for expression of islet specific markers revealed manifold enhancement of *PDX1*, *NGN3* and *PPY* genes in grafted pancreas at day 7 and 30 as compared to MSC cells. PDX1 and NGN3 showed a decline of expression after an initial rise at day 7 as expected. Enhanced PPy expression was seen by day 30.

The decellularized mouse pancreas were also recellularized with human MSC, the pancreas regenerated both islets and the exocrine pancreas. The histology of pancreas was restored and these pancreata were functional in the mice when transplanted. These cells could be additionally tracked with human nuclear antigen in the pancreas when recovered even up to 40 days (Figure 4). No adverse immune rejection was seen and the recovered pancreas stained with human nuclear antigen showed the presence of human cells also (Figure 5). The presence of human cells up to 40 days was also confirmed by using human specific Alu repeat probes on sections of pancreas recellularized with human placental MSC cells. Acinar and islet cells stained positive for the human specific Alu repeats. Pancreas recellularized with mouse adipose remained unstained with Alu repeat probes and showed reaction with mouse specific B1/B2 Sine repeats (Figure 5). There was no cross-reactivity. The rabbit pancreas recellularized with human placental cells also showed a positive reaction to Alu repeats (Suppl. Figure 2).

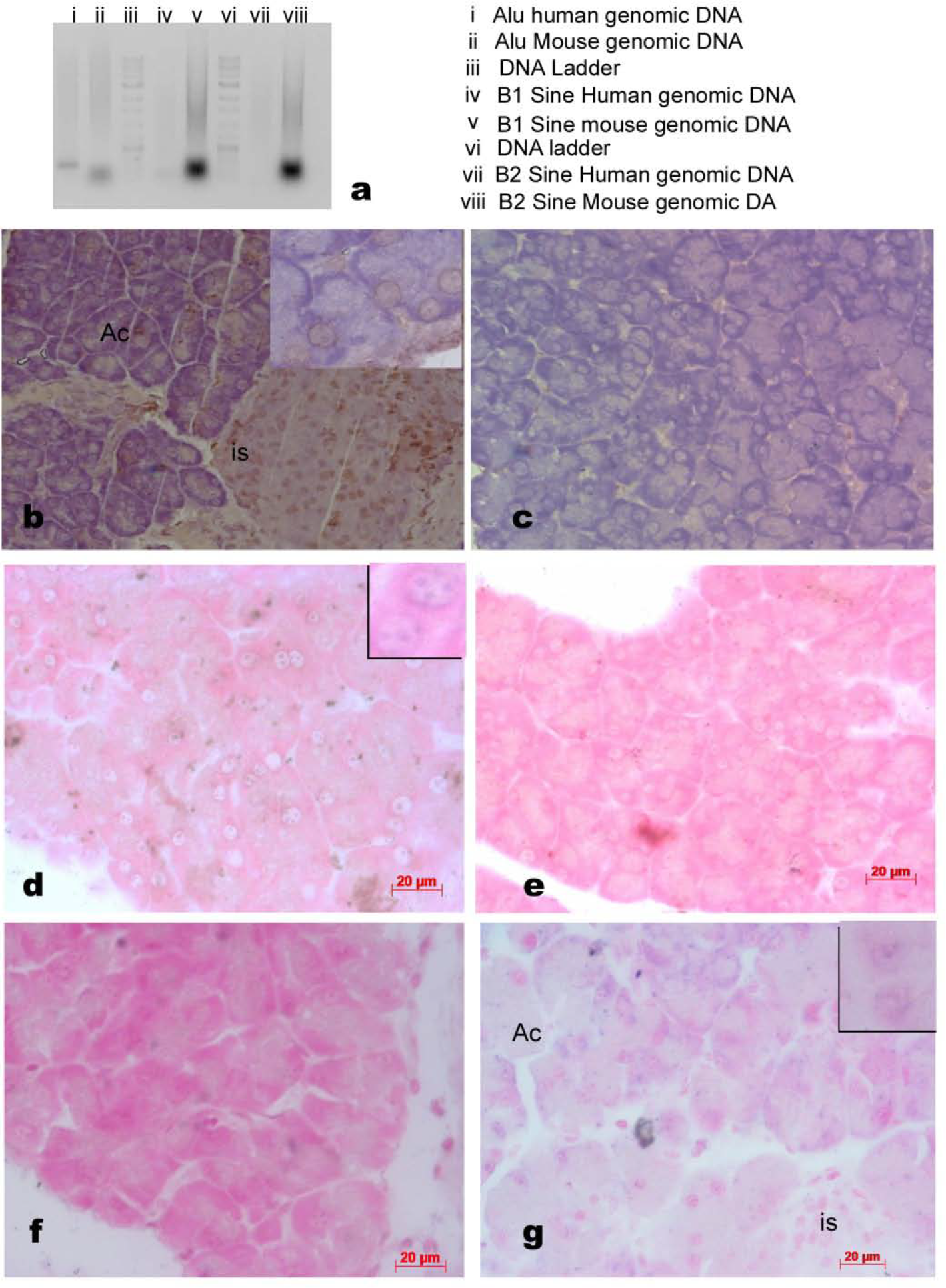

The mouse that received recellularized grafts showed regulation of glucose levels though none achieved gluconormalcy. The animals that received the recellularized pancreas with human cells started showing improved levels of human insulin and c-peptide by day 30 (Table 2). Initially, the animals continued to show reduced levels of mouse Insulin and c-peptide up to day 10. Human insulin and c-peptide were undetected at day10 and day 15 but one could detect higher levels of Human insulin and c peptide in fed animals at day 30. Pancreas recellularized with mouse adipose MSC, the animals did not show human insulin and C-peptide detection. The mouse insulin and mouse C-peptide did show improved levels.

In few attempts we had tried to graft a decellularized scaffold without recellularization but neither did the graft get cellularized nor did it help the animal regain gluco-regulation. The graft was invaded by granulocytes.

Recellularized organs were initially grafted at a subdermal location centrally on ventral side and harvested after 15 days; these mice maintained a higher level of glucose. The organ was completely invaded by granulocytes and the graft was rejected. We did not attempt grafting human or rabbit recellularized pancreas in xeno-models

## 4. Discussion

This study reports a recovery of mouse pancreas after decellularization and seeding with MSC cells, and applicability of the protocol to pancreas from rabbit and human foetus. The process of decellularization and recellularization works well for soft organs and organs with complex cell types like pancreas. Recellularization with adult mesenchymal stem cells can successfully regenerate the structure and function of the organ. Recipient animals are able to regulate blood glucose levels and restore normal histology of the organ. So far there was no immunogenicity reported when the implants were placed intra-peritoneally up to 40 days in animals.

The main considerations for the protocol were as follows (*i*) complete removal of cellular material and debris; (*ii*) removal of nucleic acid residues/components; (*iii*) preservation of ECM structure and composition; (iv) minimal loss of active components for regeneration; (*v*) functional restoration of the organ, and (*vi*) no immunogenicity.

There are a number of strategies available to decellularize organs; we chose to use continuous flow perfusion of Triton X −100, a non-ionic detergent followed by enzyme DNase to remove even traces of nucleic acids. Triton X 100 is able to disrupt lipid-lipid and lipid protein interactions, while leaving protein–protein interactions undisturbed (Seddon et al., 2004). This helps loosen the cellular material and the perfusion process per se may help in removal of the cell debris. Our protocol resulted in complete removal of cellular material as also reported by several groups (Mirmalek-Sani et al., 2013, Peloso et al., 2016). We have used the perfusion via the hepatic portal vein to get a complete decellularized pancreas. A few initial trials of perfusion via the pancreatic duct were unsuccessful as the vascular system offers better reach in the entire organ.

Proteomic analysis of the scaffolds obtained revealed the ECM proteins to be the major components left behind including Collagen type I, type VI, laminin and Proteoglycans. The immunochemistry confirmed their architectural organization. Another important constituent of the ECM - the proteoglycans, that gives the required stiffness to the matrices, were also shown to be present both in proteomic analyses and by staining of the sections. There are conflicting reports on use of Triton X-100; as it is known to remove GAGs in Aortic valve decellularization protocol (Grauss et al., 2005) and retain cell debris (Scarrit et al., 2015). Also, decellularization using Triton X-100 was shown to be most suitable for getting a clinical grade scaffold (Peloso et al., 2015, Vavken et al., 2009). Our studies confirm Triton X-100 to be suitable in all respects as the scaffolds obtained are rich in both, GAGs and matrix proteins as seen by staining and proteomics. Our decellularization protocol seems to take care of the two important considerations for obtaining an acellular scaffold; removal of cell debris and maintenance of ECM composition and structure. Triton X-100 has yielded a scaffold without any cell debris as proteomic analysis scored low number of proteins and majority of these were ECM related.

Another point of concern was nucleic acid residues left behind in the scaffold; residual DNA fragments in the scaffolds could cause immune- compatibility issues. Apart from this pathogenic viruses could pose a risk to recipients, it is better that the residues are removed by additional endonuclease treatment. The goal was to have DNA below 50ng ds DNA per scaffold and the fragment length should not be more than 200 bases (Crapo et al., 2011). Our preparations also conform to these standards, as the amount of DNA recovered per scaffold was around 50ng and the gel analysis revealed it to be fragmented.

One also looks for minimal loss of active components for growth and differentiation. The scaffold obtained is checked for its biocompatibility. The ECM components are known to support the very nature of cells residing therein (Hynes, 2009, Salvatori et al., 2014, Watt et al., 2013). Our concern was that when we seed the recovered scaffold with MSC; these MSC should be able to home in; attach and differentiate into the functional cells within the exocrine and endocrine units of pancreas. Our proteomics studies did reveal retention of PDGF like growth factors/cytokines therein and possibly the architectural/ compositional or 3D ECM cue (guides) that the stem cells infused into the pancreas were indeed able to differentiate into islets, acinar and other cell types to form a functional pancreas. We did not require any additional growth factors for the cells to differentiate into the two types of zone specific cell types. Though staining of the decellularized structure for growth factors did not show conclusive presence of growth factors (data not shown).

Pancreatic cells have great deal of plasticity inherently as shown earlier (Juhl et al., 2010, Thorel et al., 2010); where extreme loss of beta cells upon exposure to diphtheria toxin was overcome by α cells giving rise to beta cells. It was important to show that the cell implanted during the recellularization makes the pancreas functional. Human cells infused in the scaffolds could be identified by human nuclear antigen staining and by *in situ* hybridization using human specific Alu repeat probes up to 40 days; or by tracking the graft for up to 3 weeks post transplant (supp Figure 2). Scaffolds without reseeding cells, when implanted did not function alike or survive.

How many cells need to be infused was another query we addressed. Some of the scaffold recovered after just infusion of 3 × 10^5^ cells; did show cells attached to the scaffolds though the density was very less (Figure S1). A complete impregnation of the mouse scaffold with cells happened at a density of 6 × 10^5^ infused in three sets at interval of 2 hrs (Figure S1). The histology of pancreas distinctly shows the formation of islets and the acinar cells over a period of time. The histology and immuno-fluoroescence studies using antibodies against pancreatic exo- and endocrine markers clearly show distinct expression of region specific proteins. Expression of islet specific markers as estimated by qPCR was seen to increase manifold in grafted pancreas. Expression of PDX1 and NGN3 seems to follow the developmental pattern as the levels are higher in the early stages and are lower comparatively by day 30 when the pancreas seems to be histologically fully recovered. PDX1 and NGN3 show a peak in early pancreatic development and are restricted to few cell types towards maturity (Wilson et al 2003, Miyatsuka et al. 2006, Lyttle et al. 2008). PPy can be seen only in later stages when the pancreas seems to be functional by day 30. The gene expression pattern also points to *de novo* development of pancreatic cells.

One does not need to over-seed the scaffold with cells; as there may be some kind of quorum sensing for the number of cells to be retained. Quorum sensing would involve cells determining their population density due to cells engaging in neighbor communication owing to autocrine signaling (Doğaner et al., 2015). The number of cells seeded in the scaffold of rabbit and fetal human pancreas were 10 ϗ10^5^.

Grafted pancreas developed vascular anastomoses in surrounding adipose tissue in the region of implantation. Vascular net work was intact and was seen to be fully developed in histology preparations of grafted pancreas of mouse and human fetal origin. The vessels in fact can be seen loaded with blood cells in the transplanted pancreas. In rabbit recellularized pancreas, the vessels were well formed with some eosinophilic stains in it.

The functionality of constructs was confirmed in the transplantation studies where the recellularized scaffold was implanted in the peritoneal cavity near the pancreas. The streptozotocin treated mice regain gluco-regulation within a period of 25 days. Complete gluconormalization was not achieved but most of the animals started maintaining lower glucose levels compared to diabetic mice. Analysis of insulin and c peptide levels in the serum of the transplanted animals also confirmed the recovery of function as the increased serum levels of human insulin and c peptide coincide with the decline in glucose levels. Importance of ECM–cell interaction has been demonstrated by earlier studies (Nagata et al., 2001, 2002; Perfetti et al. 1996). Islet survival over the ECM and increased insulin secretion when grown in contact with ECM both *in vitro* and *in vivo* were also demonstrated in the study of De Carlo et al. (2010). ECM indeed forms a dynamic complex structure that can control cellular morphogenesis, proliferation, differentiation, lineage maintenance and ultimately function of the cells. Lineage restriction has been demonstrated when the ECM from liver was seeded with hepatocyte stem cells and kidney was seeded with ES cells (Ross et al., 2009, Wang et al., 2011). The control animals that are streptozotocin treated but did not receive the graft continue to remain hyperglycemic, substantiating functional recovery of the implanted mass. We need to improve on the period of recovery as the normalization takes about 15-20 days post-implantation. The exocrine function was not demonstrated but one can see expression of carboxypeptidase in acinar cells. Another indirect proof of exocrine activity was difficulty in RNA isolation due to the proteolytic activity.

Our studies clearly establish that extracellular matrix rich organ scaffolds prepared by decellularization is the way for creating artificial pancreas for transplantation. Upon seeding with patient specific MSCs one can expect to develop a functional pancreas. The trials with a full-fledged adult human pancreas may require preclinical standardization.

## 5. CONCLUSIONS

Pancreas derived ECM rich scaffolds can be created using Triton X based solutions. These scaffolds have all the necessary information to make a functional bioengineered pancreas using adult stem cells.

## 6. Acknowledgements

Authors wish to thank Indian council of Medical Sciences New Delhi for financial support to carry out this work. Grant no.

## Author contributions

K Uday Chandrika did all the experimental work

Rekha Tripathi standardized the decellularization protocol

Avinash Raj T helped with histology

N. Sairam helped with animal studies

Vasundhara Kamineni Parliker provided human fetal samples

Swami VB helped with proteomic studies

Nandini Rangaraj helped with confocal imaging

Mahesh Kumar J standardized animal surgeries

Shashi Singh concept, planning and writing the manuscript.

## Supplementary data

**Supplememtary Table 1.**
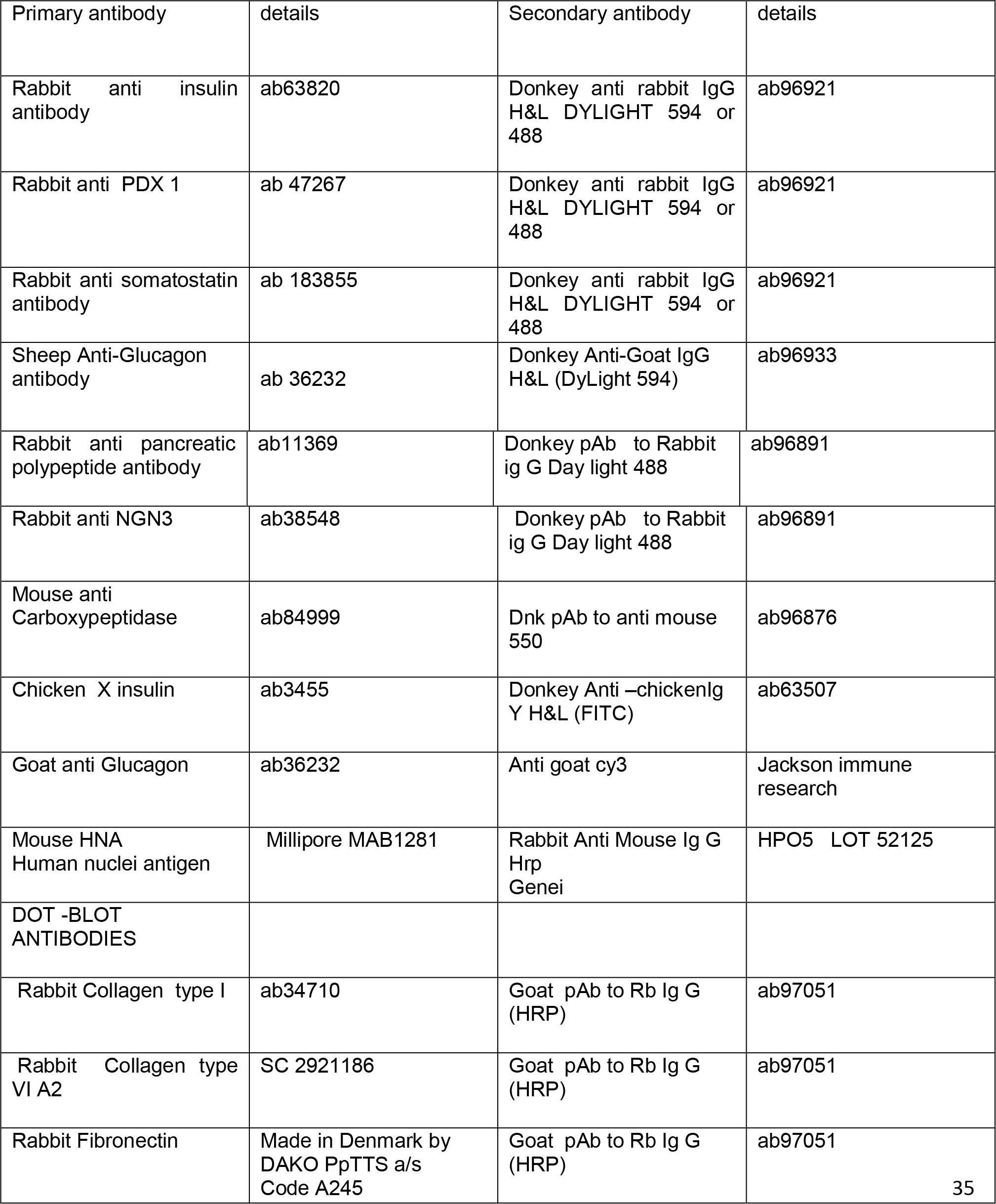
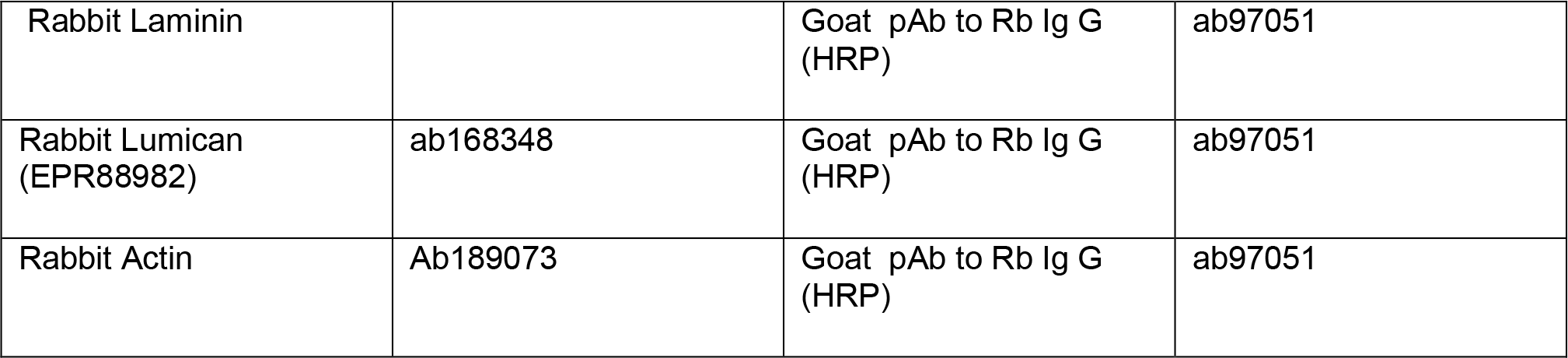
List of antibodies

**Table 2a.**
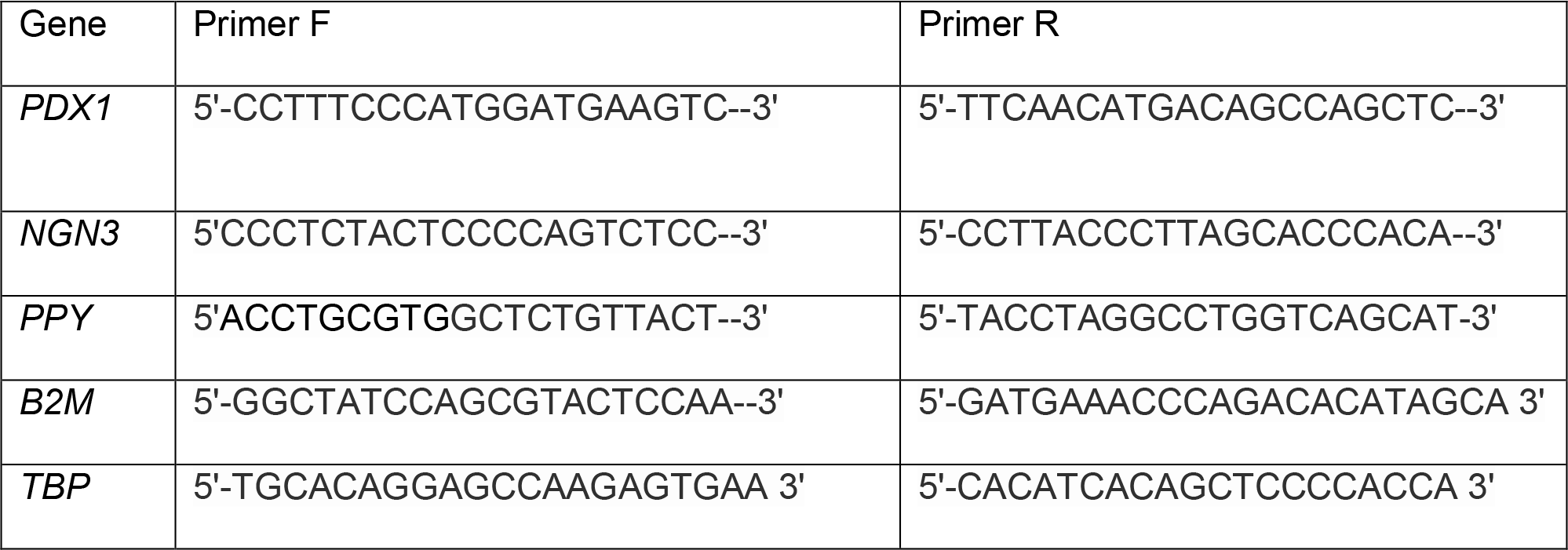
Sequence of Primers used for Real Time PCR.

**2b.**
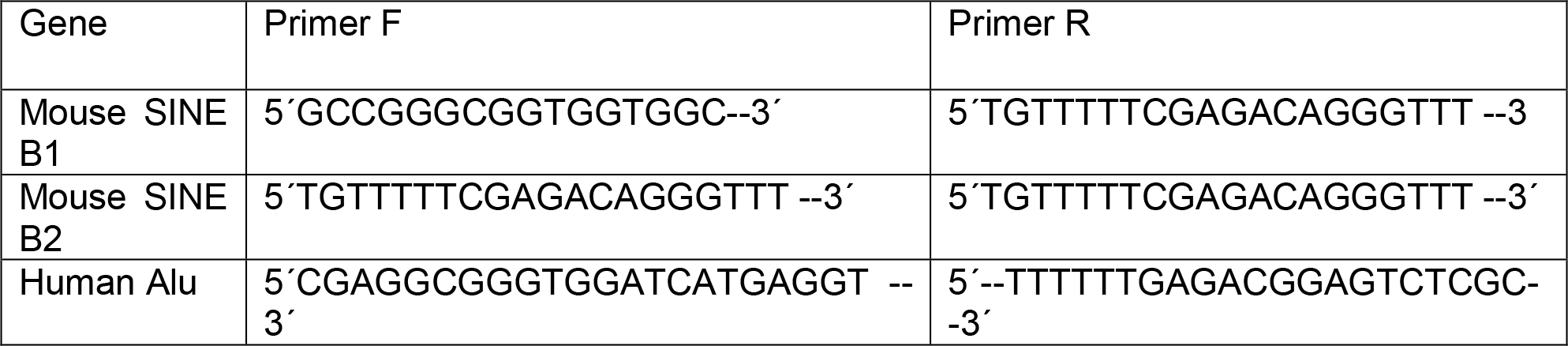
Sequence of Primers for *in Situ* Hybridization

**Figure S1.**
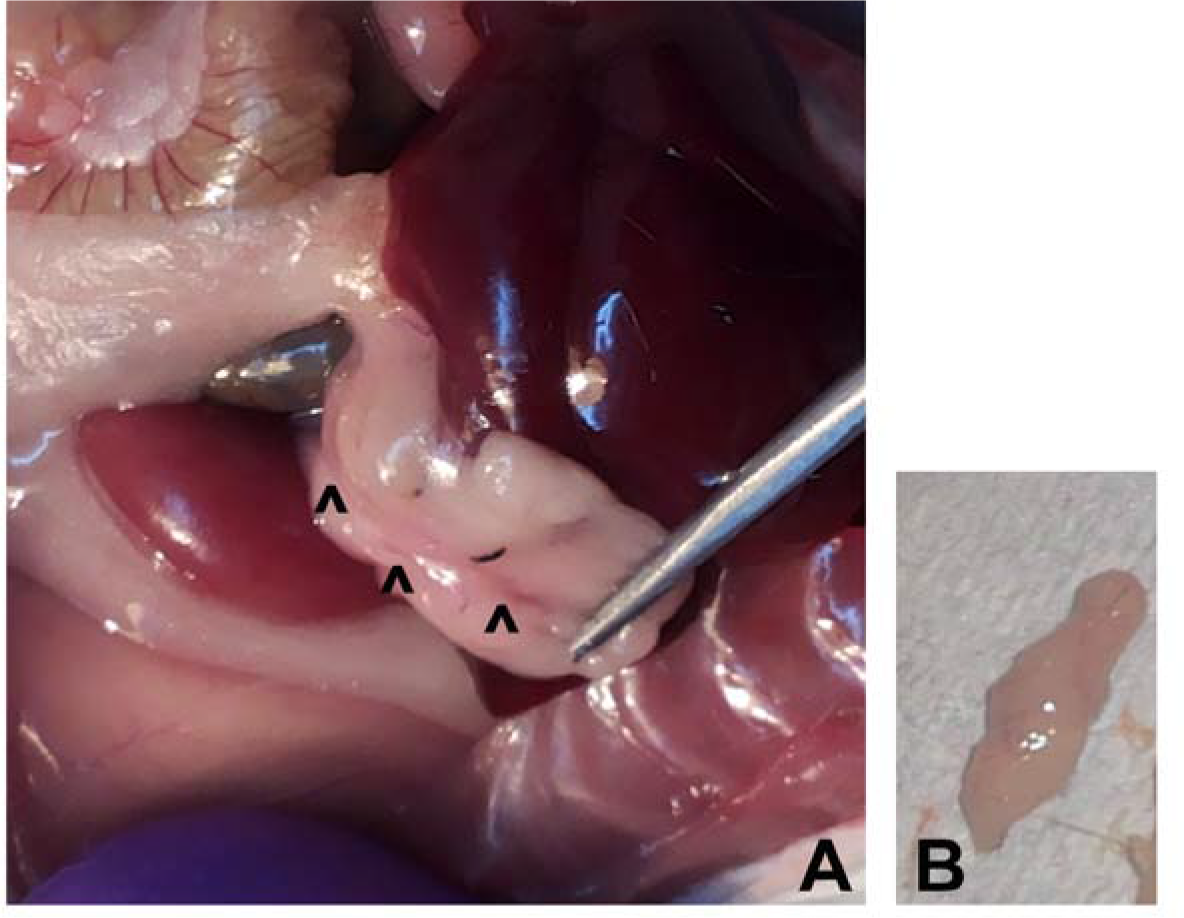
Recellularized pancreas. A. The pancreas implanted in diabetic mouse restores vasculature and vascular anastomosis (^) can be seen between the implanted pancreas and surrounding adipose. B. The recellularized pancreas excised from the animal.

**Figure S2.**
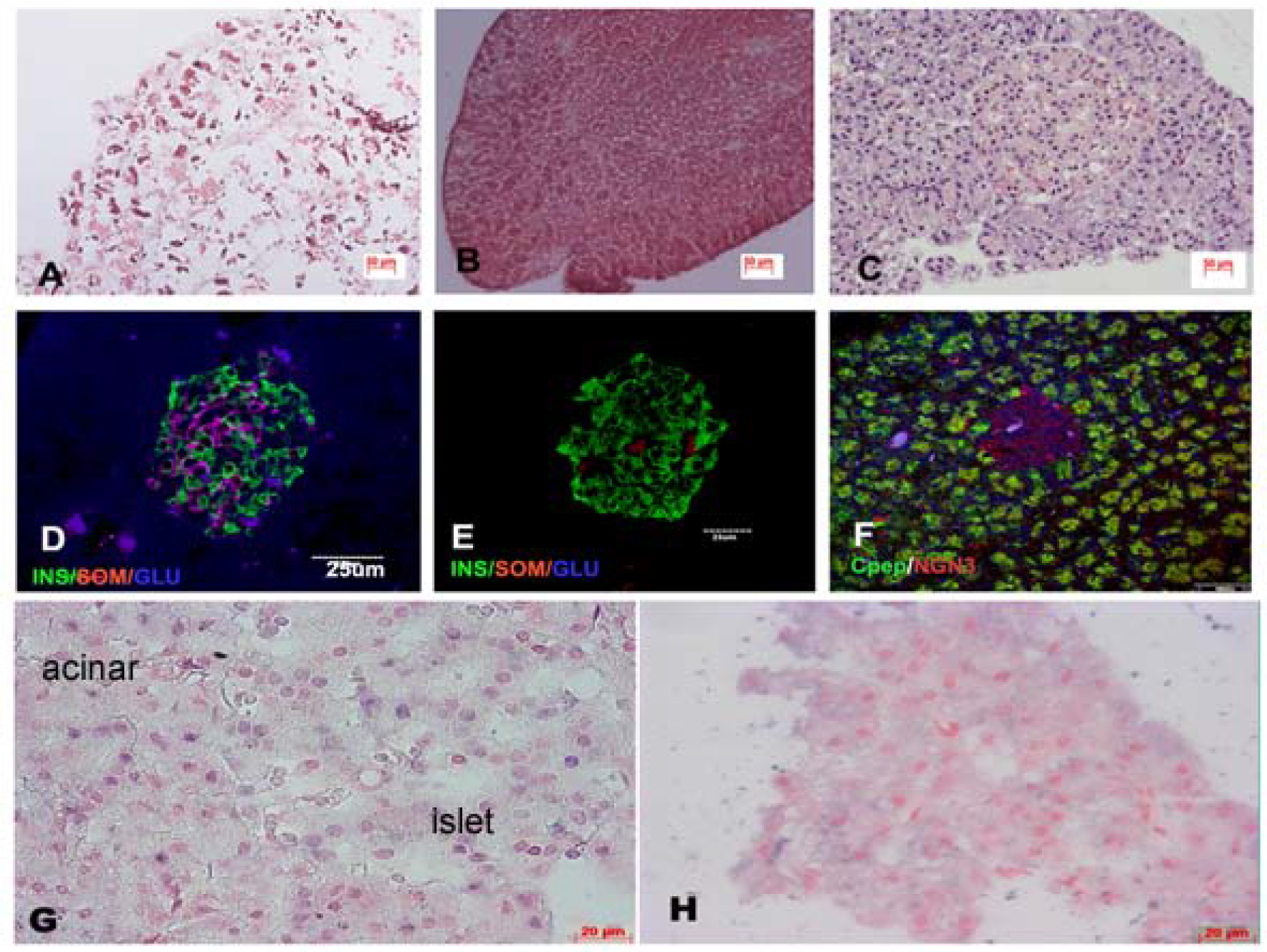
Recellularization of pancreas. Mouse pancreas recellularized with 2 × 10^5^ cells show scattered distribution and adherence of cells (A). Mouse pancreas with 6 × 10^5^ cells in fused in three rounds show better adherence and distribution of cells (B). Rabbit pancreas after recellularization with human MSC - after cells are infused the pancreas are perfused with culture medium for 10 days (C). The organ shows complete normal histology with well differentiated islets and exocrine pancreas. Immunohistochemistry of pancreas after recellularization (D-F). D is mouse pancreas infused with human MSC stained for insulin. Glucagon and somatostatin, E and F are rabbit pancreas, insulin/glucagon and somatostatin (E) and carboxypeptidase/*NGN3* (F). *In situ* hybridization of rabbit pancreas recellularized with human PL-MSC with Human Alu – repeat probes to show the differentiated cells are derived from human source (G)cells from islets and acinar region show bluish purple staining in the nuclear region; the sections when hybridized with Mouse Sine repeats B1 B2 are negative (H).

**Figure S3.**
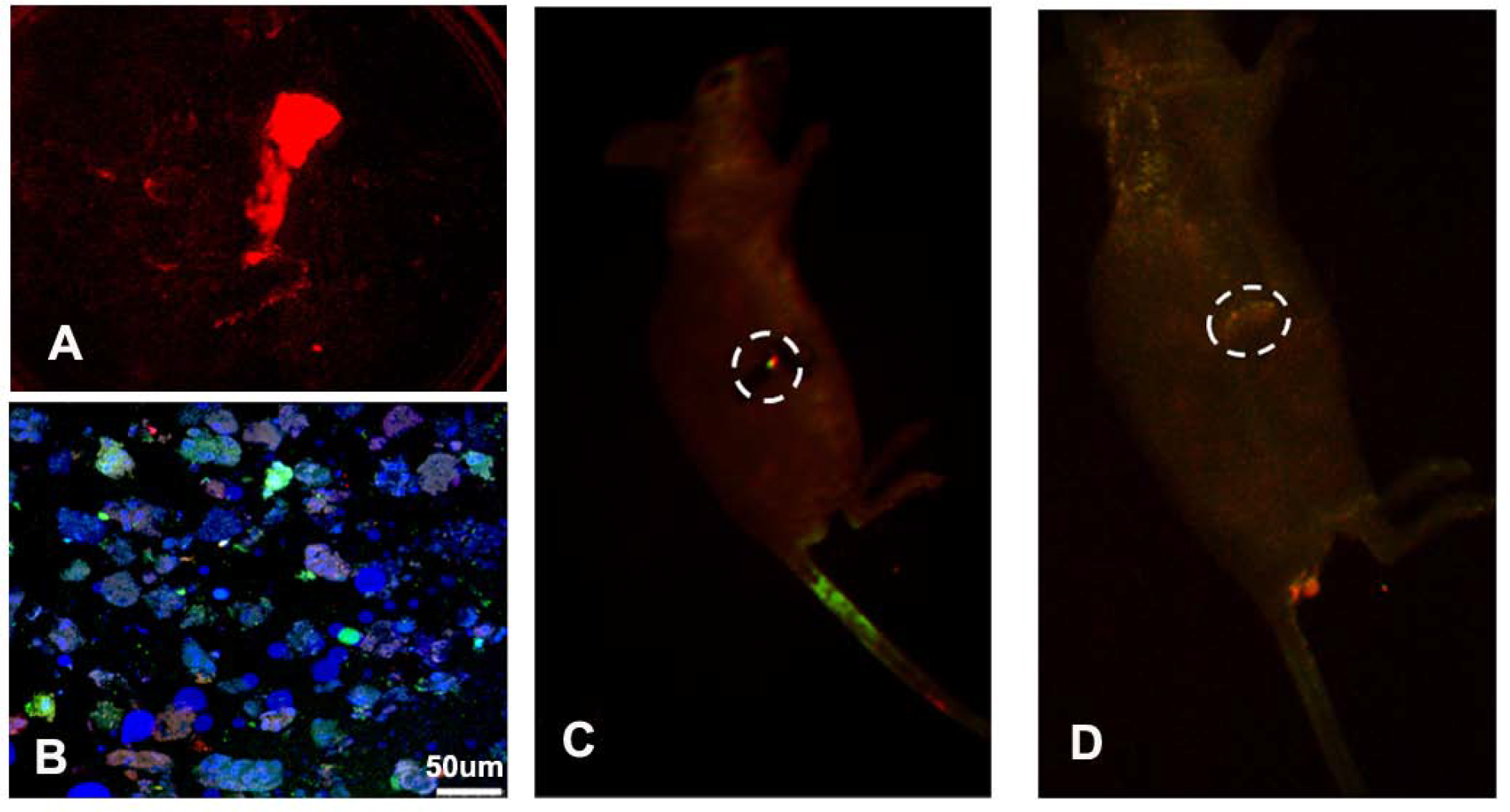
Mouse pancreas recellularized with human MSC stained with tracker dyes and implanted in nude mice. These mice were imaged in KODAK *in vivo* multispectral Imaging system Fx (Carestream Molecular imaging). The cells before infusion were stained with tracker dyes CM-Dil [Ex. 553; Em 570] (Life technologies), CFSE [Ex. 492; Em 517] (Life technologies), Cell tracker blue (CMAC) dye [Ex. 371; Em 464] (Life technologies). A. Image of pancreas before implantation. B Recellularized pancreas infused with human MSC. Cells in each round of infusion were stained with tracker dyes red, blue and green. The sections show cells stained with three dyes distributed randomly. C- *In vivo* imaging of the mouse two days after implantation shows the illuminated pancreatic region (dotted lines).D- *In vivo* imaging of the mouse after 15 days of implantation, Pancreatic region still shows fluorescence.

